# Transparent Weighting of Heterogeneous Evidence for Auditable Candidate Prioritization

**DOI:** 10.64898/2026.05.14.725271

**Authors:** Tue Nguyen Mau

## Abstract

**Motivation:** Differential-expression and network-based prioritization capture complementary molecular evidence. However, integrating these criteria poses a trade-off between score integration and methodological transparency. Existing approaches derive weights through optimization against predictive or benchmark metrics, or use manual or equal-weight settings without explicitly encoding the rationale for relative criterion importance.

**Results:** We introduce AHP-CDS, an auditable candidate-ranking method that makes criterion importance explicit through weights derived from the Analytic Hierarchy Process from explicit pairwise judgments, which allows the resulting rankings to be evaluated through consistency, sensitivity and contribution analyses. The resulting pairwise judgments achieved acceptable consistency (Consistency Ratio = 0.050). The leave-one-criterion-out analysis showed that ranking sensitivity followed the assigned weight ordering: the removal of Fold-change producing most of the disruption (Spearman compare to full method *ρ* = 0.796), whereas removing betweeness causes the least (*ρ* = 0.992). Score decomposition further made individual rank changes traceable to their evidence dimension.

**Availability:** all code is uploaded on https://github.com/NguyenMauTue/PRIDE-breast-cancer-exosome-biomarker-discovery

**Supplementary:** Is all posted online

## 1. Introduction

Triple-negative breast cancer remains clinically challenging due to substantial molecular heterogeneity (Ryu and Sohn, 2021; Senkowska et al., 2026) and limited targetable pathways (Senkowska et al., 2026). Exosome-based proteomic profiling offers a promising liquid-biopsy window into tumor biology (Ruhen and Meehan, 2019). However, translating high-dimensional measurements into a prioritized, interpretable candidate set remains difficult under small sample sizes and diverse evidence types.

Differential-expression-only prioritization can miss functionally important but moderately expressed proteins, while its network-centrality counterparts ignore signal strength; combining both has precedent in differential-topology ensembles (Pashaei et al., 2025) and expression-informed network propagation (Ren et al., 2019). Integration relocates rather than resolves the problem: criteria’s relative importance must still be specified. Existing frameworks resolve this either through: optimization against a benchmark or predictive objective (Hutz et al., 2008; Mostafavi et al., 2008; Sun et al., 2009); through unexplained manual or equal-weight settings (Xie et al., 2015; Nguyen et al., 2015), not an explicit, domain-level judgement of relative importance.

The Analytic Hierarchy Process (AHP) addresses this directly: criterion priorities are derived from pairwise comparisons, with a Consistency Ratio (CR) quantifying judgment coherence (Saaty et al., 2022). This makes the preference structure explicit and auditable, rather than a default, tuned, or optimized parameter. AHP has been previously used in gene selection (Nguyen et al., 2015), though criteria there were weighted equally to avoid subjective judgment — a limitation we address rather than sidestep.

An auditable weighting scheme characterizes only the aggregation rule. It neither explains why an individual candidate’s rank moved, nor establishes that the re-prioritization is useful: a transparent method can still fail to surface a portfolio meaningfully different from simpler baselines.

We therefore present AHP-CDS — an AHP–driven composite scoring framework that derives explicit, auditable and biologically motivated weights for multiple prioritization criteria. Using TNBC exosomal proteomics datasets as an illustrative case, we address three questions: (1) does AHP-CDS weight integration meaningfully alter prioritization relative to single-criterion and equal-weighted baselines; (2) which candidates are most affected, and through which criteria; (3) is there independent evidence that the re-prioritization is useful?

## 2. System and methods

The following sections describe the construction of the input features used by AHP-CDS. The pipeline proceeds in five stages: (i) Dataset and quality control, (ii) Missing-value handling, (iii) Differential expression analysis, (iv) pathway enrichment and candidate curation, (v) protein–protein interaction network construction with centrality calculation. The purpose of this system is to focus on the enriched pathways and generate four quantitative criteria (fold-change, statistical significant, degree and betweenness) that serve as input for the ranking algorithm.

### 2.1. Dataset and quality control

The analysis was performed on a publicly available label-free quantification dataset retrieved from PRIDE: PXD056161 (Zaripov et al., 2025), using the MaxQuant output. The experiment comprised exosomal fractions from MDA-MB-231 (TNBC) and MCF-10A (non-tumorigenic) cell lines. Each of these conditions has three biological replicates, with two technical replicates per sample.

Protein groups were retained if they had at least two unique peptides, sequence coverage *≥* 5%, were not a reverse hit, potential contaminant, or identified solely by a modification site. LFQ intensities were extracted, with zero values treated as NA and log2-transformed prior to downstream analysis. A protein was considered detected in a biological replicate if its aggregated intensity was non-missing.

Technical replicates were assessed using pairwise correlation and principal component analysis. No technical replicates were excluded. Technical replicates were aggregated to the biological-replicate level by taking the mean of non-missing values (or the single available value when one technical replicate was missing). Proteins were retained for downstream analysis when detected in at least two of three biological replicates in at least one condition.

### 2.2. Missing-value handling

Missingness was modeled as abundance-dependent via logistic regression against protein intensity using technical-replicate observations. The observed abundance-dependent detection pattern suggested imputation using a left-censored Gaussian approach following Perseus strategy (Tyanova et al., 2016), with a downshift of 1.8 standard deviations and a width of 0.3. Imputation parameters were estimated independently per group rather than pooled across the dataset, motivated by the low inter-sample variability observed within PXD056161 itself (coefficient of variation *≈* 0.71%, median/MAD basis, across the six biological-replicate samples), which supported treating each condition’s intensity distribution as sufficiently homogeneous to estimate its own shift/width parameters without borrowing information across groups. The imputed dataset was used for downstream differential expression analysis.

### 2.3. Differential expression

Differential expression analysis was performed using `limma` (Ritchie et al., 2015). A design matrix without an intercept (*∼* 0 group) was fitted, with the contrast defined as MDA-MB-231 minus MCF10A. Protein-level statistics were estimated using `lmFit, contrasts.fit`, and empirical Bayes moderation with eBayes using default `limma::eBayes` setting, followed by `topTable`. P-value were adjusted for multiple testing using the Benjamini–Hochberg procedure to obtain false discovery rates (FDRs). Rather than applying a hard threshold, FDR and logFC was retained as a continuous scoring criteria within the composite scoring framework.

### 2.4. Pathway enrichment and candidate curation

Proteins were ranked by moderated t-statistic and analyzed using gene set enrichment with gsePathway from ReactomePA. UniProt identifiers were mapped to Entrez IDs; when multiple proteins mapped to the same Entrez ID, the protein with largest absolute t-statistic were retained. Enrichment analysis was performed with `minGSSize` = 10, `maxGSSize` = 500, `pvalueCutoff` = 0.05, following the normalized enrichment score (NES) formulation of GSEA (Subramanian et al., 2005). Pathways were retained using criterion of a supplementary magnitude filter of |NES| *>* 1.5. This magnitude threshold is not part of the original GSEA significance criterion (which specifies FDR *q <* 0.25 on its own) and is instead an author-chosen filter intended to focus on pathways with a stronger effect size.

Candidate curation was restricted to core-enrichment proteins belonging to the retained enriched pathways, thereby focusing the candidate universe on biological processes showing evidence of phenotype–associated dysregulation. The resulting candidates were mapped to HGNC symbols and UniProt Swiss– Prot identifiers using `Ensembl BioMart` and annotated using Gene Ontology terms. Candidates were then filtered to remove common contaminant or housekeeping/cytoskeletal proteins, including histones, keratins, actins, and tubulins (Welsh et al., 2024), together with a manually curated black list of six additional genes (Supplementary ).

### 2.5. Protein–protein interaction network and centrality metrics

A protein–protein interaction network was constructed using the STRING REST API from the curated candidate gene set. Gene symbols were resolved to STRING identifiers for *Homo sapiens*, and interactions with a combined STRING confidence score *≥* 0.7 were retained. The resulting undirected network was restricted to candidate set, such that centrality measures reflected the relevant topological position of candidates within the phenotype-associated candidate space. Duplicate edges, self-loops, nodes without retained interactions were excluded. Raw degree and betweenness centrality were calculated using igraph

## 3. Algorithm

### 3.1. AHP–weighted Composite Decision Score

AHP–CDS is a multi-criteria scoring framework that aggregates *n* quantitative criteria into a single prioritization score for each candidate. AHP is used to derive weights (Saaty et al., 2022). The procedure consists of four steps.

1. Pairwise comparisons among the criteria are elicited on the Saaty scale and assembled into a reciprocal judgement matrix A
2. Criterion weights **w** are obtained by the standard approximate eigenvector method (column normalization followed by row averaging). Consistency of the judgments is quantified by the Consistency Ratio, which is regarded as acceptable if *≤* 0.1 (Saaty et al., 2022)
3. Each criterion is transformed (if required by its distribution) and scaled to a common range, typically [0, 1], before weighting. The specific transformation is context-dependent and left to the user.
4. The CDS of candidate *i* is the weighted sum

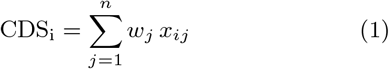

where *x*_*ij*_ is the normalized value of criterion *j* for candidates *i*.

## 4. Implementation

### 4.1. Data and code availability

Proteomic data were retrieved from PRIDE (PXD056161). External database queries (STRING REST API, Reactome via ReactomePA, Ensembl BioMart, and the Comparative Toxicogenomics Database) were all performed in August 2026; because STRING and CTD are continuously updated resources, results reflect the state of these databases at the time of query rather than a fixed release. All analyses were performed in R; full package version information (sessionInfo()) and the complete analysis pipeline are available on GitHub at https://github.com/NguyenMauTue/PRIDE-breast-cancer-exosome-biomarker-discovery.

### 4.2. Case-study parameterization

In this study, we incorporated four criteria: absolute FC, negative FDR, degree, and betweenness centrality. Absolute FC was log-transformed into |log_2_ FC|, FDR into *−* log_10_ FDR, whereas the two centrality measures were log(1+x)–transformed, then min-max normalized. Relative importance was *a priori* on biological and statistical grounds: FC received the highest weight as a large abundance difference is essential for experimental follow-up in noisy exosomal liquid-biopsies; degree was ranked next as a measure of local network importance within the candidates; FDR was down-weighted because of the limited statistical power of the n = 3 design; betweenness was assigned the lowest weight because of it is a global topology measure that depends on shortest-path structure and can therefore be sensitive to incomplete or study-biased interaction coverage in aggregated PPI networks (Blumenthal et al., 2024; Salehzadeh-Yazdi and Hütt, 2025). The weights are treated as expert judgment rather than parameters to be optimized against a ground-truth, thus no data-driven re-weighting was performed. The resulting weights were 0.518, 0.164, 0.242, and 0.076 for absolute FC, FDR, degree, and betweenness, respectively, with a Consistency Ratio of 0.050.

### 4.3. Benchmarking methods

To assess the effect of AHP–derived weighting, we compared AHP–CDS against three internal baselines that operate on the identical normalized criterion matrix:

- **FC-only**: ranking by log FC;
- **Centrality-only**: ranking by degree centrality;
- **Equal-weight**: the same four criteria combined with uniform weights (*w*_*j*_ = 0.25), reflecting the equal-weighting strategy previously adopted in AHP gene selection to avoid subjective judgement (Nguyen et al., 2015)

These ablations isolate the contribution of the AHP weighting scheme from the contribution of multi-criteria integration itself. As an external comparator, we implemented a faithful algorithmic analogue diffusion-based ranking following the Global Ranking procedure of Ren et al. (2019). The algorithm uses a random-walk normalized Laplacian and a discrete heat-kernel approximation (*N* = 3, *α* = 0.5). Three deliberate adaptions were made to the original protocol: (i) the interaction network was taken from STRING (combined score *>* 0.7) and binarized, compared to HINT interactome; (ii) the seed set was defined as proteins significant under Bonferroni correction of limma p-values, preserving a single differential-expression pipeline across the study; (iii) the diffusion truncation parameters were kept exactly as published. Candidates absent from the resulting network received the worst rank. The differential–topology method DiCE (Pashaei et al., 2025) was considered but not included, because its phenotype-specific edge re-weighting requires Pearson correlations estimated from only three samples per condition.

### 4.4. Evaluation procedures

The evaluation framework was designed to assess AHP– CDS from three complementary perspectives. Rank–level evaluation examines whether AHP–derived weighting produces a ranking that differs or agrees with alternative prioritization strategies. Criterion-level evaluation assesses the contribution and sensitivity of individual evidence dimensions, determining whether the resulting prioritization depends disproportionaly on any single criterion. Portfolio-level evaluation examines the composition and balance among the high ranked candidates, addressing whether the ranking produces a biologically informative candidate set. Together, these analyses distinguish ranking agreement, criterion dependece, and candidate-set utility.

Agreement between AHP–CDS and each internal baseline was quantified by Spearman rank correlation, Kendall’s *τ*, and Rank-Biased Overlap (RBO; Webber et al. (2010)) at pre-specified depths. Per-candidate rank displacement (Δ*r*) and top-*k* membership changes were also recorded.

For each candidate *i* and criterion *j*, the additive contribution *C*_*ij*_ = *w*_*j*_*x*_*ij*_ was computed. Leave-one-criterion-out (LOCO) sensitivity analysis was performed by sequentially removing each criterion and recalculating the composite ranking to assess the dependence of candidate prioritization on individual evidence dimensions (Doğan, 2026).

Candidates were assigned to evidence tiers on the basis of curated gene–disease associations retrieved from the Comparative Toxicogenomics Database (Davis et al., 2025): Tier 1 (breast cancer association), Tier 2/3 (other cancer association or significant network neighbor of Tier 1), and Tier 4 (no supporting association). This reflects a gradient of candidates’ evidentiary directness: from expected in the context of breast cancer, to having suggestive yet not confirmed disease evidence, to novel prioritization. Portfolio balance at depth *k* was summarized by the evenness index

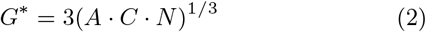

where *A, C* and *N* are respectively the proportions of Tier-1, Tier-2/3, and Tier-4 within top*k. G*^*\**^ quantifies whether a ranked candidate set retains representation across the predefined evidence tiers. One consequence of the geometric-mean construction is that G* is invariant to which of the three terms carries given value, so two top-*k* sets with permuted (*A, C, N* ) triples yield an exact same *G*^*\**^. *G\** is designed to capture a method’s capacity to simultaneously recover known biology, extend evidence-linked reasoning to plausible new candidates, and surface understudied signal within a single ranked output. The geometric mean equals zero whenever any component is empty: a zero in Tier-1 indicates that the selected set contains no positive controls; in Tier-2/3, a lack of evidence-linked rationale bridging findings to confirmed biology; or in Tier-4, that it does not extend prioritization into candidates lacking supporting disease associations. However, one consequence of the geometric-mean construction is that G* is invariant to which of the three terms carries given value. Observed values were placed in the context of the distribution obtained by repeated random sampling of *k* candidates.

## 5. Results and Discussion

We used the pipeline described in 2 to PXD056161 and yielded a final candidate pool of 143 proteins. This pool reflects successive scoping filters: retention within pathways passing an enrichment magnitude and statistical significance threshold, restriction to the leading-edge proteins driving that enrichment within each retained pathway, and having at least one high-confidence STRING interaction within resulting candidate subnetwork itself. Therefore, the pool concentrates on proteins that are both leading contributors to strongly altered pathways and topologically connected to one another.

The methodological rationale in 2 establishes how heterogeneous evidence can be integrated through an explicit weighting scheme. The relevant question for the results is therefore what the resulting weighting changes in candidate prioritization. We examine whether AHP–CDS produces a materially different ranking from single-criterion and equal-weighted alternatives, which criteria account for the largest changes in individual candidates, and whether the resulting re-prioritization provides independent evidence for biologically plausible candidates. Throughout the analyses below, we report outcomes at two selection depths: *k* = 15 as the primary depth and *k* = 10 as the secondary check at a narrower selection window.

**Table 1.** AHP Pairwise Comparison Matrix and Derived Criterion Weights. Judgments follow the Saaty (1980) scale: 1 = equal importance, 3 = moderate, 5 = strong, 7 = very strong, 9 = extreme; reciprocal values indicate inverse preference. Consistency Ratio (CR) = 0.0503 *<* 0.10, confirming acceptable judgment consistency. FDR was down-weighted relative to network degree centrality (0.1639 vs. 0.2419) to reflect the limited statistical power inherent to small-*n* proteomics datasets (*n* = 3 per group), wherein FDR-adjusted *p*-values may be overly conservative and less discriminative than topological network evidence.

| Criterion | FC | Degree | Betweenness | Weight |  |
| --- | --- | --- | --- | --- | --- |
| FC | 1 | 4 | 2.5 | 5 | 0.5181 |
| FDR | 1/4 | 1 | 1 | 2 | 0.1639 |
| Degree | 1/2.5 | 1 | 1 | 5 | 0.2419 |
| Betweenness | 1/5 | 1/2 | 1/5 | 1 | 0.0761 |

### 5.1. Rank-level evaluation

Table 2 reports rank-level agreement between AHP-CDS and each internal baseline across the 143-candidate pool. Concordance is highest against Equal-weight (Spearman *ρ* = 0.904, Kendall *τ* = 0.736), and lowest against Centrality-only (*ρ* = 0.602, *τ* = 0.454), leaving FC-only intermediate (*ρ* = 0.876, *τ* = 0.684). Despite the near-identical overall correlation with Equal-weight, individual-candidate displacement remains substantial: 71.3% of candidates move by more than 5 ranks, and 27.3% move by more than 20 ranks, with a maximum single-candidate shift of 42 positions. This indicates that even with the criterion set held fixed, the AHP-derived weighting itself materially reshapes individual rankings. Divergence from Centrality-only is more pronounced across every metrics: 84.7% of candidates shift by more than 5 ranks and 49.7% by more than 20 ranks, with a maximum displacement of 86 positions. Rank-biased overlap at depth 10 (RBO = 0.399) indicates that only a minority of the top 10 ranking structure is shared between the two methods.

Top-k membership changes this pattern. At *k* = 10, AHP-CDS shares 8, 5, 2 out of 10 with FC-only, Equal-weight, Centrality-only, while at *k* = 15, FC-only and Equal-weight converge to the same overlap (11/15). Centrality-only remains lowest at 7 out of 15. This suggests that FC being the highest-weighted criterion (0.518), dominates specifically at the top of the distribution, where extreme FC values determine membership regardless of the other three criteria. While AHP-CDS ranking across full candidate pool are more similar to a genuine multi-criteria composite. The two metrics therefore are not in tension, they characterize different regions of the ranking. A distinction examined directly through per-criterion contribution shares in Section 5.2.

### 5.2. Criterion-level evaluation

Contribution-share distribution explains the top-k/whole-pool inversion noted in Section 5.1: FC’s mean proportional contribution is higher among the top-15 candidates than among the remaining pool (*P*_FC_ = 0.531 *>* 0.416), while degree and betweenness revert the pattern (*P*_Deg_ = 0.216 *<* 0.278, *P*_Bet_ = 0.058 *<* 0.087). Meanwhile, FDR remains comparatively stable in both depths. This is consistent with the shape of each criterion’s normalized distribution rather than from the weighting itself: the implied mean normalized value is the lowest for FC (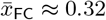, consistent with the right-skewed distribution of FC magnitudes), while being highest for FDR 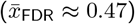. Consistent with this, FC is the only criterion whose pool-average realized contribution 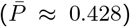 is lower than its assigned weight (*w*_FC_ = 0.518). FC’s high weight translates into outsized influence specifically for the FC–extreme minority at the top of the ranking, while contributing comparatively less across the rest of the pool.

Leave-one-criterion-out (LOCO) sensitivity analysis confirms the weight ordering directly: removing FC causes the largest disruption (Spearman *ρ* = 0.796 versus the full ranking, 80.4% of candidates shift by ¿5 ranks; max Δ*r* = 87), followed by degree (*ρ* = 0.947, 64.3%), FDR (*ρ* = 0.981, 41.3%) and betweenness (*ρ* = 0.992, top-10/15 membership unchanged). This matches the ordinal AHP weight ranking with (FC = 0.518, FDR = 0.164, Deg = 0.242, Bet = 0.076). The dependence is also direction-specific, tracking the criterion each baseline omits at T = 15: vs. FC-only, promoted candidates show higher *P*_Deg_ (mean *P*_Deg_ = 0.415 vs. 0.097); vs. Centrality-only, promoted candidates show higher *P*_FC_ (mean *P*_FC_ = 0.601 vs. 0.253 demoted), a similar pattern holds for T = 10. Each single-criterion baseline is corrected by AHP-CDS in the direction traceable to the criterion it omits.

### 5.3. Portfolio-level evaluation

We examine portfolio composition at four selection depths (*k* = 5, 10, 15 and 20). These depths mark the transition in how the methods stand relative to a null reference band derived from 10,000 random draws of size k from the 143-candidates pool. At *k* = 5, RWR reaches the null 95% upper bound (*G*^*\**^ = 0.952) while AHP–CDS remains below it (0.865). At *k* = 10, AHP– CDS ties the bound at 0.991, whereas at *k* = 15, AHP–CDS reaches its peak at 1 and falls back to 0.991 at *k* = 20 exceeds the bound.

Decomposing *G*^*\**^ into tier counts shows that the two external and internal comparators diverge from AHP–CDS for different reasons. Of the 18 Tier-1 genes in the pool, AHP–CDS accumulates them steadily with depths (3, 4, 5, 6 at four k milestones). By contrast, RWR under this projection recovers exactly two Tier-1 candidates at every depths; the additional candidates it ranks between *k* = 5 and *k* = 20 are drawn entirely from Tier-2/3 or Tier-4. However, in this setting, RWR’s *G*^*\**^ never exceeds the null band. This pattern is best interpreted as the portfolio composition obtained when a ranking generated under a different candidate definition is evaluated in the present universe, rather than as evidence that random–walk diffusion is inherently unable to enrich for curated positive controls.

Equal-weight recovers Tier-1 candidates at parity or above AHP–CDS (for example, 4 vs. 3 at *k* = 5 and 6 vs. 5 at *k* = 15). Yet, its *G*^*\**^ never exceeds the null band. The reason is structural: at k = 5, Equal-weight’s top 5 contains no Tier-4 candidates, collapsing the three-way evenness index across all three evidence tiers. AHP-CDS diverges from RWR primarily in Tier-1 recovery, whereas it diverges from Equal-weight in maintaining three-way balance.

These comparisons are purely exploratory. The reference distribution is constructed over a small and tier-imbalanced pool. This leads to limited resolution in the reference distribution, particularly at small k. No correction for multiple depths was applied. AHP–CDS surpasses the null band at two depths only (*k* = 15 and 20). The portfolio-balance advantage is therefore scale-dependent, appearing most clearly at intermediate selection depths. From this perspective, AHP– CDS produces a different profile of tier coverage: balancing from positive control (Tier-1), under–annotated (Tier-4), and evidence-linked rationale bridging findings to confirmed biology candidates (Tier-2/3).

### 5.4. Case-study

The seven case studies illustrate how the composite score behaves when different evidence dimensions agree, compete, or remain absent. Rather than treating the AHP–CDS rank as a self-sufficient verdict, we decomposed the score to examine which components determine each candidate’s position. Visualization of the rank changes and LOCO without FC (see mega figure 2) revealed two recurring patterns: the direction of movement between the FC-only and Centrality-only baselines, and the direction of rank change after removing FC from the composite. We therefore use these two observed patterns to organize the case studies, with the former describing baseline rank movement and the latter identifying the criterion that rescues the composite rank.

**Figure 1.**
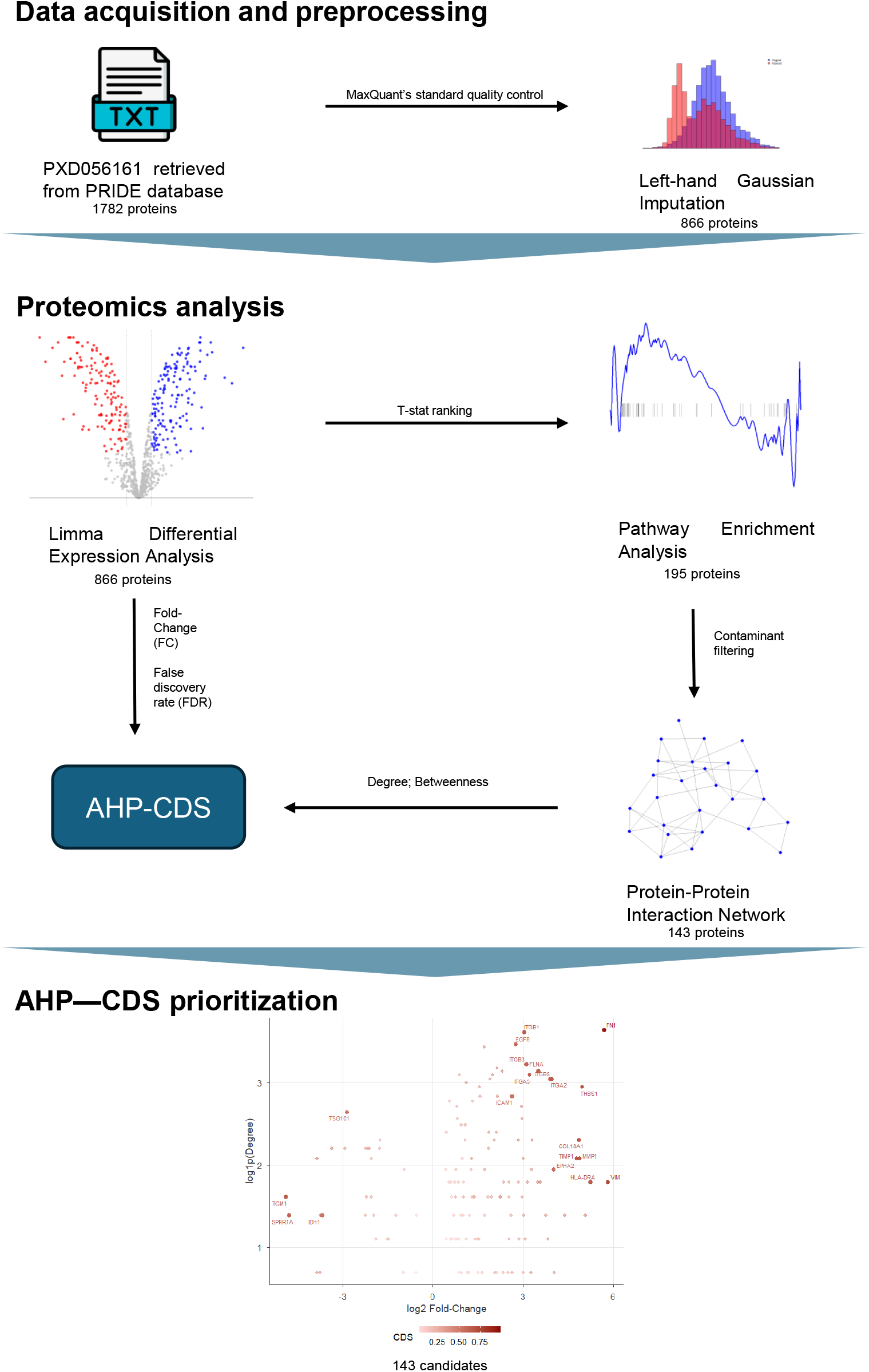
Pipeline Overview. Multi-stage screening and candidate attrition workflow: starting from 1,782 raw identified proteins, 866 proteins passed quality control and low-abundance filtering for differential expression analysis (limma). Gene set enrichment analysis identified 195 core-enrichment proteins within enriched pathways (|NES| *>* 1.5). Filtering out contaminants and blacklisted genes yielded 143 candidate proteins, which were mapped to construct a PPI network for topological analysis. Multi-criteria prioritization framework. Four quantitative metrics—Fold-Change (FC) and False Discovery Rate (FDR) from differential expression, together with Degree and Betweenness centralities from network analysis, were extracted for the 143 candidates and integrated into the AHP-CDS algorithm to produce the final prioritized target list.

**Figure 2.**
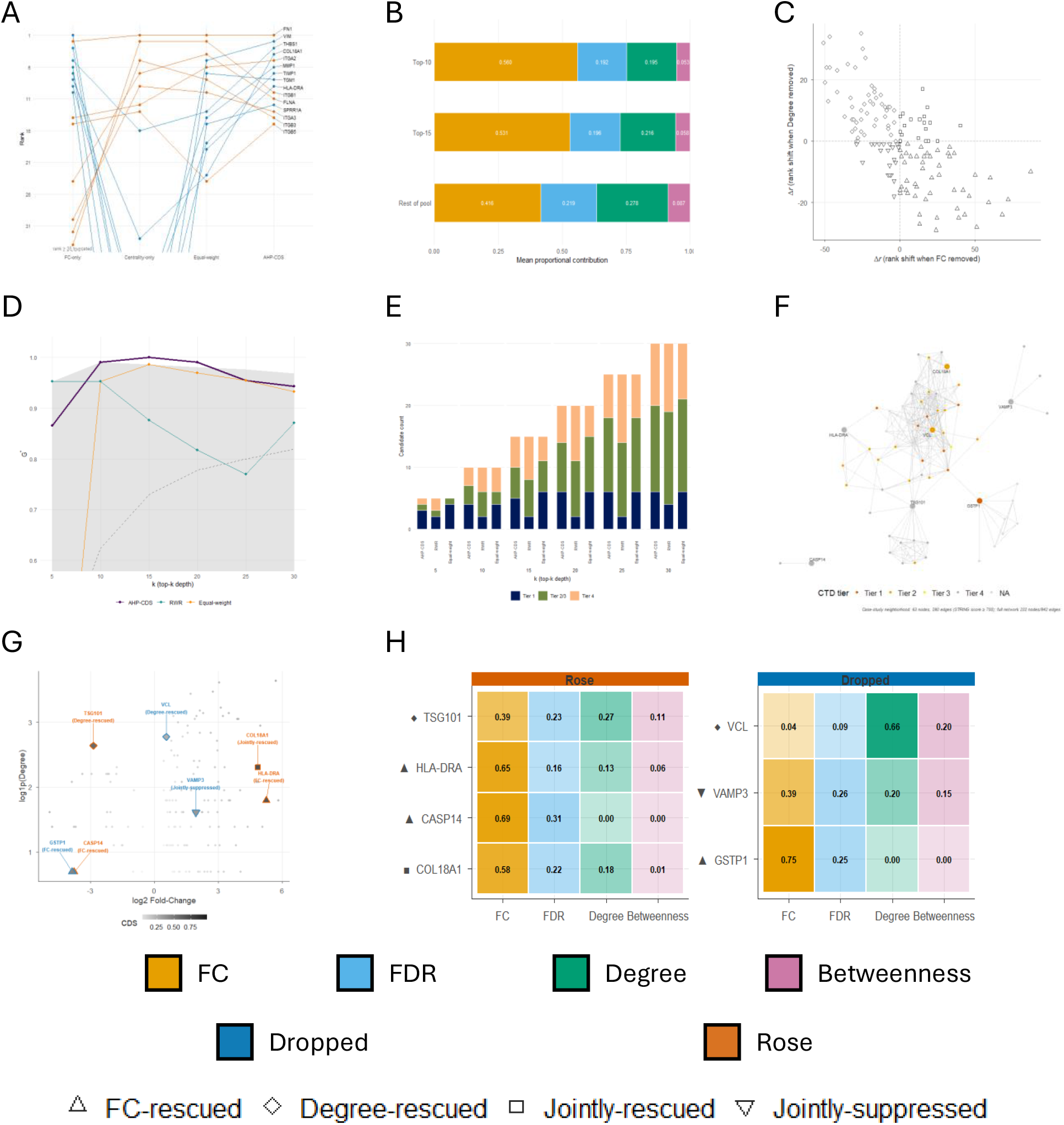
Multi-layer evaluation of the AHP-CDS ranking framework. Shared legend consists of colors: FC=gold, FDR=light blue, Degree=teal, Betweenness=magenta (B,H); AHP-CDS=crimson, RWR=grey-blue, Equal-weight=tan (D,E); Dropped (FC-favored)=blue, Rose (topology-favored)=orange (A,F,H). Shapes: *△* FC-rescued, ♢ Degree-rescued, □ Jointly-rescued, ▽ Jointly-suppressed (C,G). (A) Rank bump plot, top-15 candidates across FC-only/Centrality-only/Equal-weight/AHP-CDS. (B) Mean contribution share (FC/FDR/Degree/Betweenness) by rank tier (Top-10/15/rest-of-pool). (C) Pool-wide LOCO Δrank: FC-removed vs. Degree-removed. (D) *G*^*\**^ vs. top-*k* depth for AHP-CDS/RWR/Equal-weight, with randomized-draw null band. (E) CTD-tier counts (1/2*·*3/4) vs. top-*k* depth, per method. (F) STRING neighborhood of case-study genes; node color = CTD tier, size *∝* degree. (G) log2FC vs. log1p(Degree), shaded by CDS score; case-study candidates labeled by archetype. (H) Contribution-share heatmaps: Rose (TSG101, HLA-DRA, CASP14, COL18A1) vs. Dropped (VCL, VAMP3, GSTP1).

*GSTP1* (Tier-1) carries strong FC and FDR evidence, but no retained interactions in the candidates STRING subnetwork (*C*_Deg_ = *C*_Bet_ = 0). Its score is therefore driven entirely by expression-based evidence (*C*_FC_ = 0.329, *C*_FDR_ = 0.109), yielding rank 50 — well below its FC-only rank of 17, yet above its Equal-weight rank of 84. The drop relative to FC-only is not a failure to recover a strong signal; it reflects the cost of incorporating evidence dimensions for which the candidate has little or no supporting signal.

*VCL* illustrates the converse trade-off: strong topological support is held back by the weak FC evidence, so composite rank falls drastically compared to Centrality-only (95 vs. 22). Removing FC from the composite score (LOCO) improves its rank to 44 — a 51 position improvement, largest in the whole pool. This confirms that FC is the dominant suppressing criterion for this candidate. The case makes visible a property easily obscured by a single numerical score: multi–criteria integration necessarily creates trade-offs whose magnitude is governed by the stated weights.

*CASP14* (Tier-4, zero network contribution) is again, dominated by FC (*C*_FC_ = 0.320) and ranks 40 under AHP–CDS versus 126 under Centrality-only. Unlike GSTP1, the absence of curated cancer evidence does not preclude independent biological support; *CASP14* has been reported as upregulated in TNBC tissue (Makuch-Kocka et al., 2025; Handa et al., 2017). The weighting scheme therefore preserves a strong expression signal when complementary layers are sparse, without implying equal support across all dimensions.

*HLA-DRA* shows when the same mechanism becomes practically consequential. Also a Tier-4 and poorly ranked by Centrality-only (62), its strong expression signal places it third under FC-only and ninth under AHP-CDS. Independent TNBC evidence supports the biological relevance of this signal. Here the composite score does not merely illustrate FC weighting; it positions a candidate with incomplete curated evidence inside the top-10 of the prioritization pool.

*TSG101* demonstrates a distinct form of integration: neither single-criterion baseline alone surfaces it, yet the composite does. As a Tier-4 candidate, its expression signal alone yields only a middling: FC-only rank of 43, and its network centrality alone offers no rescue either (Centrality-only rank 25). However, the composite score places it at rank 17. Unlike *GSTP1, CASP14*, and *HLA-DRA*, where network contribution is effectively absent (*C*_Deg_ *≈* 0), *TSG101* carries a genuine, non-negligible network component (*C*_Deg_ = 0.160) alongside FC (*C*_FC_ = 0.233). As a canonical ESCRT-I subunit and standard exosome-biogenesis marker, *TSG101* carries no curated disease association in CTD whatsoever — the starkest curation gap among the candidates discussed here — yet independent evidence links its overexpression to mammary tumorigenesis and exosome secretion (Dennaoui et al., 2025; Oh et al., 2007).

*VAMP3* is reciprocal discordant case, in which the criterion contributing most to the composite score is not the criterion sustaining its rank. FC-only and Centrality-only placed it respectively at 74 and 79, whereas for AHP–CDS, it’s 64. FC contributes the largest share of composite score. However, removing FC shifts its rank by Δ*r* = *−*13, identifying FC as the criterion that suppressing its AHP–CDS rank. Therefore, *CHMP5* represents a discordant case, illustrating why the effect of a criterion within the composite cannot be inferred directly from the corresponding single-criterion baseline.

*COL18A1* provides the clearest convergence of the evidence dimensions represented in the scoring model. Ranked fourth by AHP–CDS, it improves on its already strong FC-only position (rank 8). The score retains a dominant FC contribution (*C*_FC_ = 0.426, *C*_FDR_ = 0.164) while also receiving decent network support (*C*_Deg_ = 0.132). LOCO identifies FC as the largest individual driver, yet the dependence is weaker than for candidates whose ranks rest almost entirely on expression. Therefore, *COL18A1* is not simply lifted up by the strongest criterion; its expression signal is accompanied by corroborating network evidence. Its Tier-2 assignment further distinguishes curated database annotation from broader biological evidence: although CTD records only a general cancer association, independent studies link *COL18A1* directly to breast cancer and TNBC biology (Devarajan et al., 2023).

Taken together, the seven cases show that the composite rank should neither be accepted without inspection nor discarded in favor of its components. *GSTP1* demonstrates the cost of additional evidence dimensions; *VCL* exposes the resulting tension;*CASP14* shows that an explicit weighting scheme can retain signal when other layers are absent; *HLA-DRA* shows when that mechanism becomes practically consequential; *TSG101* shows that distributed, moderate evidence across dimensions can elevate a candidate beyond what any single criterion achieves alone; and *COL18A1* demonstrates convergence rather than competition. The value of AHP–CDS does not lie in rendering any single evidence type obsolete. It lies in providing an explicit, consistency-checked rule that allows heterogeneous evidence to contribute to a common prioritization decision while preserving an auditable link between assigned preferences, criterion-level contributions, and the resulting rank.

### 5.5. Limitation

While the AHP–CDS framework enables transparent and auditable repriorization of candidates from high-throughput proteomics screening, our findings should be interpreted in the context of standard bioinformatic caveats (e.g., sample size constraints, imputation artifacts, STRING network attrition, and absence of wet-lab validation).

AHP–CDS’s Saaty weights (*w*_FC_ = 0.518, *w*_FDR_ = 0.164, *w*_Deg_ = 0.242, *w*_Bet_ = 0.076) were elicited as fixed *a priori* judgment and were not varied in primary analysis. We treat AHP as deliberate expert judgment rather than an optimization target (*CR* = 0.05 confirms internal consistency of the elicited matrix, not correctness of its content). This leaves untested how the specific pairwise-comparison magnitudes affect the resulting composite. For example, a stated 3-fold versus 5 or 7-fold preference between adjacent criteria might yield a materially different composite. Rankings should be read as reflecting one internally consistent expert judgment, not as invariant to the granularity of that judgment.

Independence between paired criteria, implicit in treating AHP–CDS as a weighted sum, was not formally tested, including for non-linear dependence. The LOCO disruption asymmetry reported in 5.2 (removing Degree causes substantially larger rank disruption than removing Betweenness, *ρ* = 0.947 *vs*. 0.992) offers indirect, partial evidence against full redundancy within each axis, but does not establish independence, and normalization choices (log 1*p* - compression for Degree and Betweenness, *−* log_10_ for FDR) further reshape each criterion’s effective contribution independently of its assigned weight. Because this pattern may be specific to this dataset’s expression and network structure, users applying AHP–CDS elsewhere shall assess pairwise dependence directly rather than assume it from this study.

Weights were elicited from a single domain-informed judgment rather than aggregated across independent raters using established group-AHP consensus methods (e.g, Saaty’s geometric mean aggregation; Saaty (1989)). Internal consistency (*CR <* 0.1) confirms the judgment is self-coherent, not that it would replicate across raters. Rankings should accordingly be read as one expert’s explicit, auditable prioritization scheme, not a consensus judgment.

AHP–CDS’s criterion set reflects one deliberate choice of evidence to integrate; the framework does not require these specific four. Incorporating additional or alternative criteria encoding qualitatively different signal, for example, an exosome-cargo propensity score reflecting secretory-pathway/ESCRT-associated packaging likelihood, or a cross-cohort reproducibility signal independent of this study’s expression data, was not attempted here and is left as a direction for extending AHP-CDS beyond this study.

## Abbreviations

TNBC: Triple-negative breast cancer
AHP: Analytic Hierarchy Process
CDS: Composite Decision Score

